# A dynamic Campylobacterales epibiont community associated with the bone eating worm *Osedax*

**DOI:** 10.1101/2022.11.14.516544

**Authors:** Shana Goffredi, Balig Panossian, Camille Brzechffa, Naomi Field, Chad King, Giacomo Moggioli, Greg W. Rouse, José M. Martín-Durán, Lee Henry

## Abstract

*Osedax*, the deep-sea annelid found at sunken whalefalls, is known to host Oceanospirillales bacterial endosymbionts intracellularly in specialized roots, that help it feed exclusively on vertebrate bones. Past studies, however, have also made mention of external bacteria on their trunks. During a 14-year study, we reveal a dynamic, yet persistent, succession of Campylobacterales integrated into the epidermis of *Osedax*, that change over time as the whale carcass degrades on the sea floor. The Campylobacterales associated with seven species of *Osedax*, which comprise 67% of the bacterial community on the trunk, are initially dominated by the genus *Arcobacter* (at early time points < 24 months), the *Sulfurospirillum* at intermediate stages (~ 50 months), and the *Sulfurimonas* at later stages (>140 months) of whale carcass decomposition. Metagenome analysis of the epibiont metabolic capabilities suggests a transition from heterotrophy to autotrophy along the successional gradient, and differences in their capacity to metabolize oxygen, carbon, nitrogen, and sulfur. Compared to free living relatives, the *Osedax* epibionts were highly enriched in transposable elements, implicating genetic exchange on the host surface, and contained numerous secretions systems with eukaryotic-like protein domains, suggesting a long evolutionary history with these enigmatic, yet widely distributed deep-sea worms

## Introduction

Whalefalls create a unique environment for deep-sea organisms as the decaying carcass serves as a bountiful, albeit ephemeral, source of nutrition on the seafloor. *Osedax* “bone-eating worms” specialize in these habitats by infiltrating and degrading the whalebones using a unique root-like tissue that contains obligate intracellular symbionts within the Oceanospirillales (1–3). This symbiosis has a profound influence on accelerating the degradation of marine mammal skeletons, and therefore nutrient remineralization and ecosystem longevity in the deep-sea (4). While many studies have examined the diversity, genomics, and physiology of the primary intracellular symbiont of the nearly 30 known species of *Osedax* (ex. 5-9), much less is known about other bacteria, including the Campylobacterales, which have been regularly recovered from the external surface of these important residents of deep-sea whalefall ecosystems (2, 10, 11).

Campylobacterota, formerly known as Epsilon-proteobacteria, are known to oxidize sulfide and other intermediate sulfur compounds and have an affinity for habitats rich in both organics and sulfides, such as hydrothermal vents, methane seeps, and whalefalls (12–14). They are now recognized as important players in deep-sea biogeochemical cycles (15–17). At whalefalls, in particular, the Campylobacterales can represent up to ~30% of bacterial ribotypes recovered from bone surfaces or sediments, compared with < 2% community membership for sediments beyond the influence of the whale carcass (18–19). It is, however, currently unclear whether the Campylobacterales found on *Osedax* are non-specific transient associations, or persistent epibionts of the worm itself.

A remarkable diversity of bacteria form non-transient associations with eukaryotes, both internally and externally, and can contribute to the health, physiology, behavior and ecology of their hosts. Bacteria that interact with surface epithelia can play important ecological roles for animal hosts by reducing exposure to harmful compounds, and modulating interactions with predators or pathogens, to name a few (20–21). The physical and chemical properties of a host surface, prevailing conditions of the surrounding seawater (in the case of marine epibionts), as well as interactions among the microbial residents, can all shape this community. Due to their different ecologies, surface-associated bacteria are often metabolically distinct from their free-living populations, demonstrating higher enzymatic activity, growth and reproduction, and increased lateral gene transfer compared to free-living cells (20, 22). Here, we present a 14-year study of the bacterial communities associated with the external surfaces of seven species of *Osedax* worms. Using molecular, metagenomic, and microscopy analyses we reveal a dynamic community of Campylobacterales epibionts associated with *Osedax* that are unique from close relatives and metabolically suited to different successional stages of whale decomposition.

## Results

To characterize the *Osedax*-associated bacterial diversity, we performed 16S rRNA gene amplicon sequencing of 37 specimens collected from two Pacific Ocean sites (Table 1; Figure 1). Based on this analysis, the Campylobacterales was identified as the dominant bacterial group associated specifically with the *Osedax* trunk (67 ± 19 %; Figure S1). *Arcobacter, Sulfurospirillum* and *Sulfurimonas* were the primary *Osedax*-associated Campylobacterales genera recovered, and specific ribotypes were distinct from those known to associate with other animals from reducing habitats (only 82% 16S rRNA gene similarity; Figure 2, Supp Materials). The only other common bacterial 16S rRNA gene barcode was from an uncultured member of the Kordiimonadales (Alphaproteobacteria; comprising 29 ± 17 % of the microbial community; Figure S1), was an uncultured member of the Kordiimonadales (Alphaprotoebacteria), closely related to those recovered previously from the external surface of *Osedax* (11) and sunken wood (58).

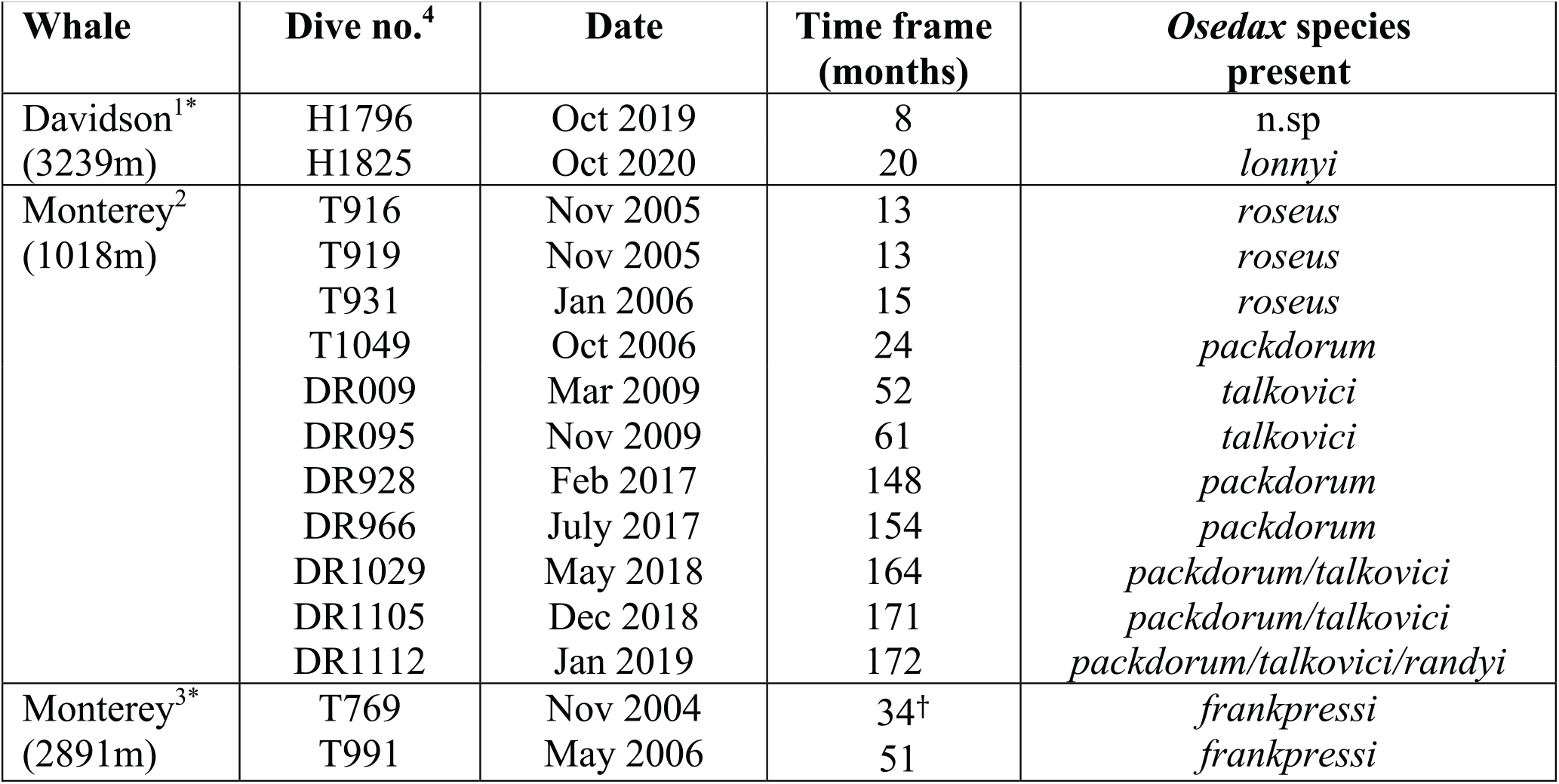

**Figure 1:**
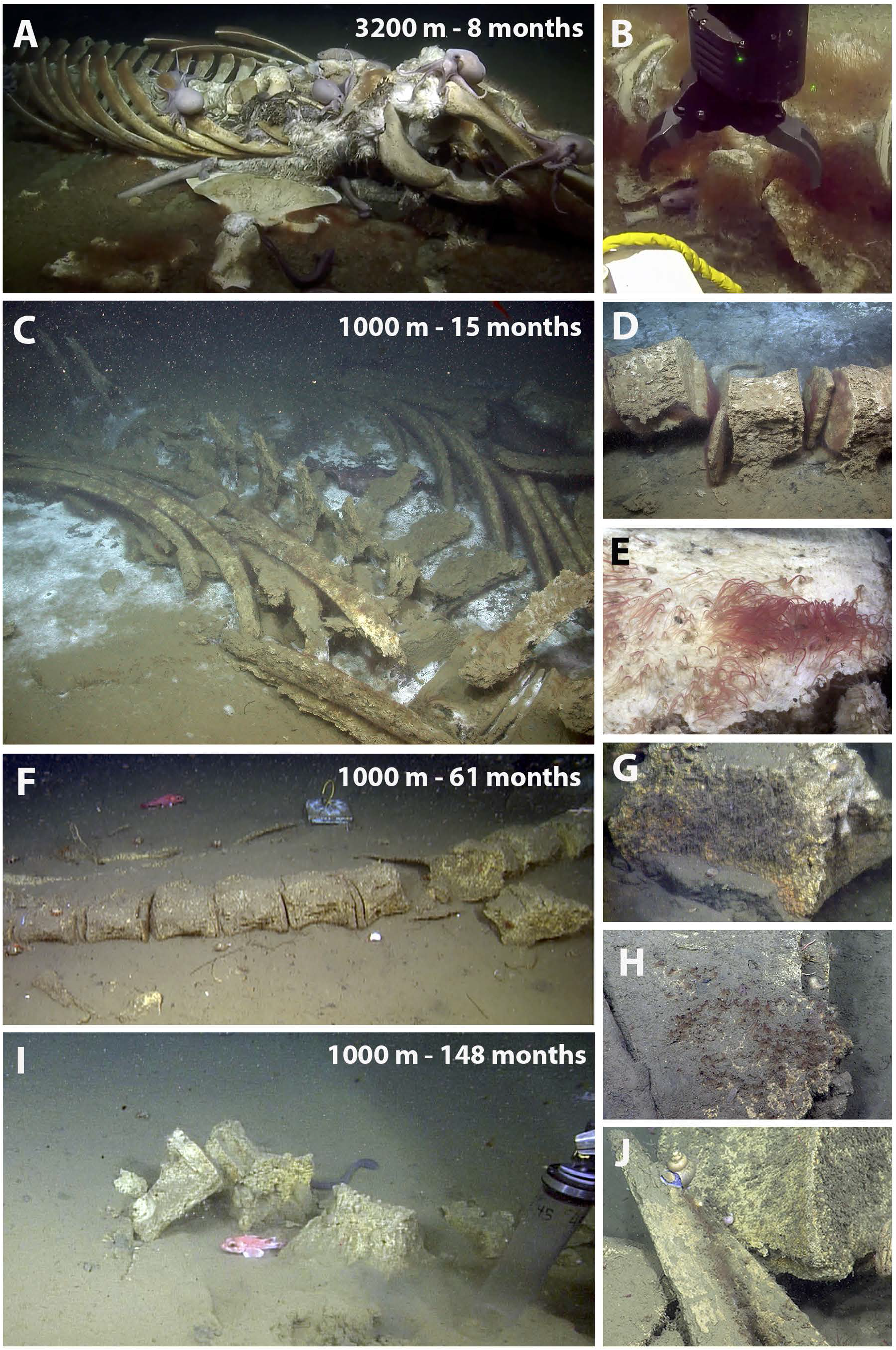
Still images of whalefalls off of northern California, USA, showing decomposition over time and condition of the carcasses at the time of sampling. A 3239m whalefall at (A-B) ~8 months (dive H1796, 10/16/19, initial observation). A 1018m whalefall at (C-E) 13-15 months since deposition on the seafloor (dives T916 and T931, 11/7/05 and 1/4/06, resp); (F-G) 61 months (dive DR095, 11/18/09); (H-I) 148 months (dive DR928, 2/23/17); and (J) 172 months (dive DR1112, 1/7/19). H = ROV Hercules, T = ROV Tiburon, DR = ROV Doc Ricketts.

**Figure 2:**
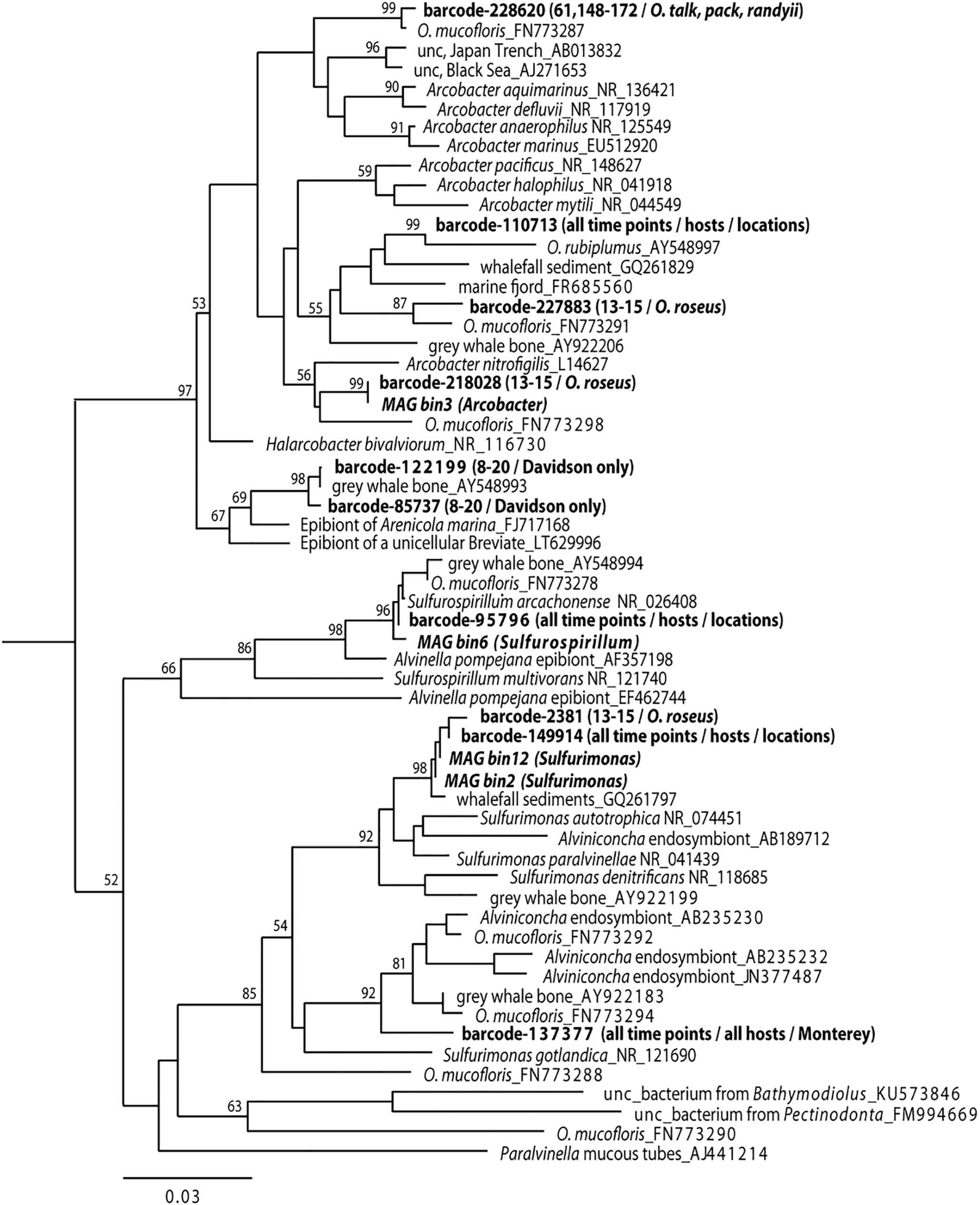
Phylogenetic relationships among the *Osedax* Campylobacterales epibionts, based on the 16S rRNA gene. Taxa in bold denote those generated in this study. *Helicobacter ganmani* (NR_024836) was used as the outgroup (not shown). Numbers at nodes indicate bootstrap support (1000 replicates, neighbor-joining, Tamura-Nei model), aligned using Geneious Prime 2021.2.2. Additional sequences from cultured representatives were obtained from GenBank, as were sequences from Suzuki et al. 2006; Goffredi & Orphan 2010; Verna et al. 2010).

Fluorescence microscopy showed a close association of the Campylobacterales with the trunk epithelial surface of *Osedax*. The Campylobacterales were the only obvious bacteria present along the epidermis, based on overlap between the universal and specific bacterial probes (Figures 3–4). They occurred along the full length of the trunk, and appeared very closely attached to the apical end of exposed epidermal labia, although some also appeared in epidermal cavities, as was seen via TEM (Figures 4C-D, S2). Slight autofluorescence of a matrix in which the bacteria were embedded, made the determination of their specific position inconclusive via FISH microscopy. For the mucous tube, which is secreted by numerous glands on the trunk and is used by the worm to glide up and down, only non-Campylobacterales bacteria were identified via microscopy (Figure S3). No bacteria were observed on plume or root surfaces (Figure 3A,G), despite significant surface area of both tissues.

**Figure 3:**
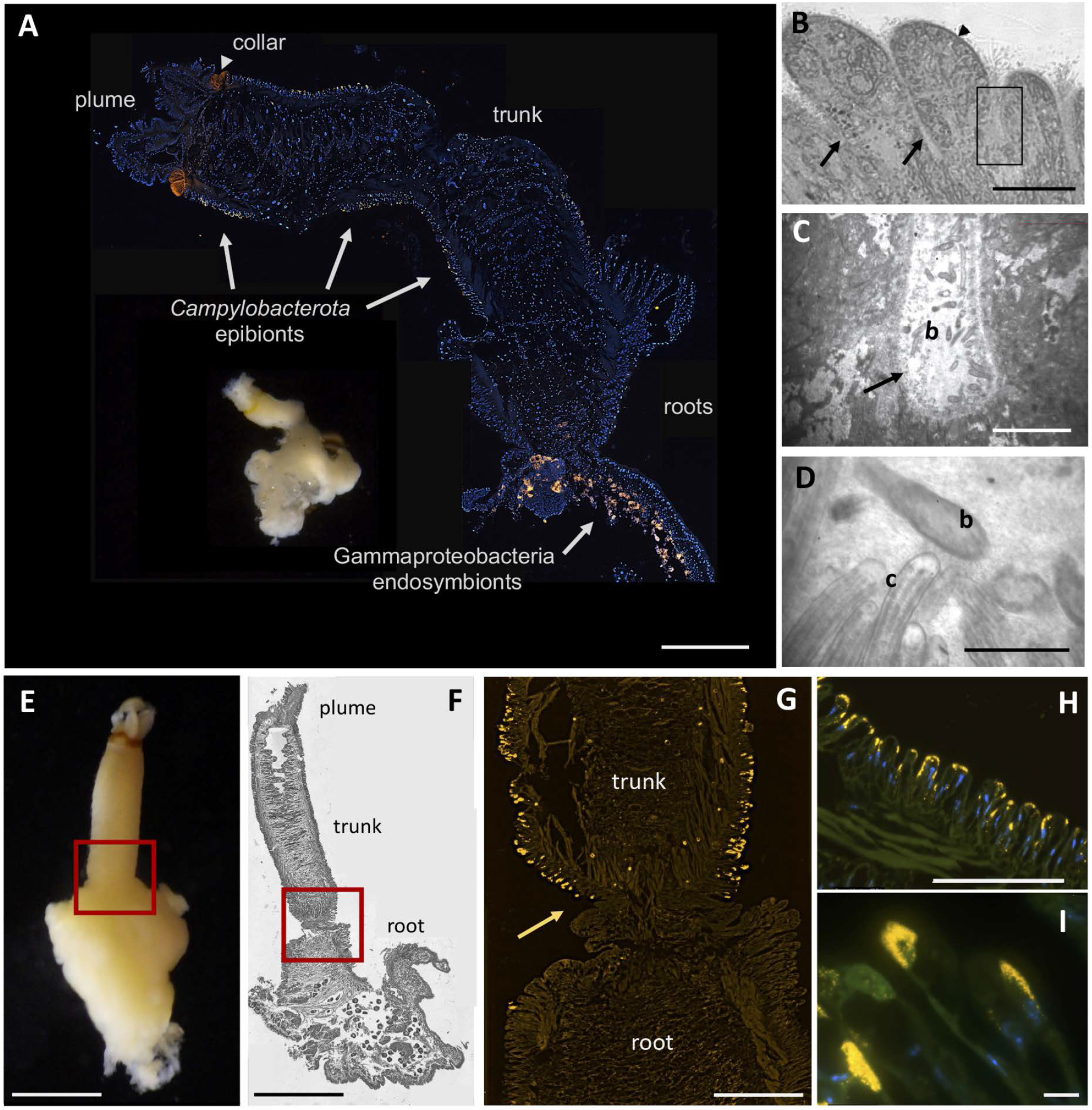
Microscopy of *Osedax packardorum*. (A) Whole image of a specimen from dive DR966, side-by-side with a longitudinal cross section, from plume to roots, hybridized with a fluorescent probe targeting all bacteria (Eub338_Cy3, shown in orange), and counterstained with DAPI, showing host cell nuclei in blue. (B-D) Transmission electron (TEM) microscopy of *Osedax* trunk tissue (dive DR1112), revealing bacteria-like cells (b) in epidermal grooves (arrows), in contact with host cilia (c, arrowhead). Square in B highlights regions in C and D. (E) Whole image of specimen from dive DR1105. (F) Light microscopy of 5-μm Wright-stained section embedded in Steedman’s resin. (G) FISH microscopy using a probe designed specifically to target *Sulfurimonas*, Epsi126_Cy3, showing the abrupt delineation in epibiont presence between the trunk and roots, at arrow, with slight autofluorescence. (H-I) FISH microscopy using probes Epsi126_Cy3, EPS549_FITC and Eub338_Cy5. Complete overlap between the probes is shown in yellow, in addition to DAPI-stained host cell nuclei in blue, with slight autofluorescence in the FITC channel. Scale bars: A, 200μm (not including inset). B, 100μm. C, 25μm. D, 1μm. E, 500μm. F, 300μm. G, 160μm. H, 100μm. I, 10μm.

**Figure 4:**
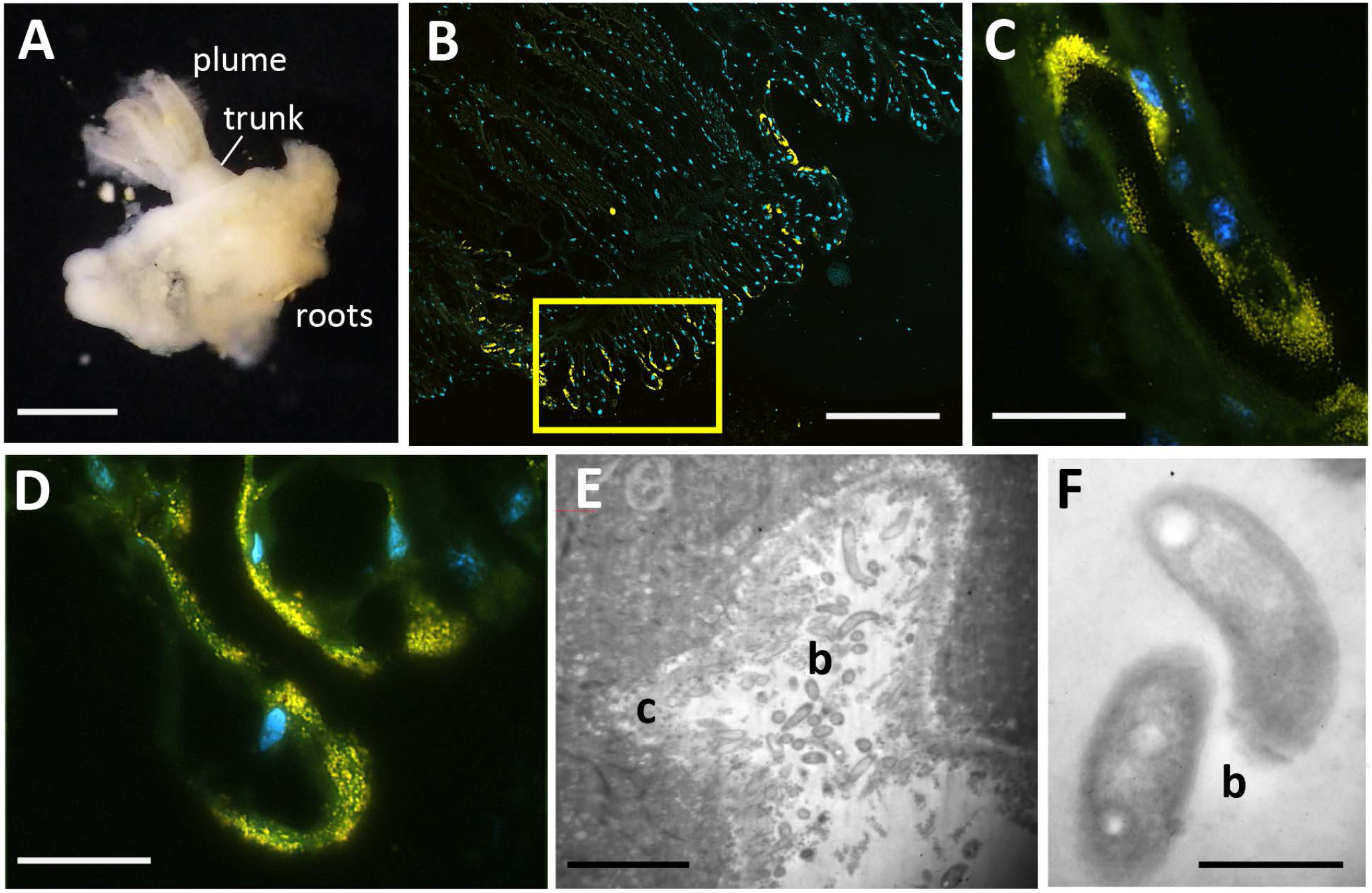
Microscopy of *Osedax talkovici*. (A) Whole image of a specimen from dive DR1112. (B-D) Fluorescent in situ hybridization (FISH) microscopy using probes Eub338_Alexa488 and EPS549_Cy3. Complete overlap between the probes is shown in yellow, in addition to DAPI-stained host cell nuclei in blue. (E-F) Transmission electron (TEM) microscopy of *O. talkovici* trunk tissue, revealing bacteria-like cells (b) in epidermal grooves, in contact with host cilia (c). Scale bars: A, 500μm. B, 100μm. C-D, 10μm. E, 20μm. F, 0.5μm.

To explore whether *Osedax’s* epibiont community changes as the whale carcass degrades, we collected the annelids from two whalefalls between 8 to 172 months (>14 years) since carcass deposition; one intentionally deposited on the seafloor at 1018m depth in the Monterey Canyon (4) and a second discovered in an early stage of decomposition at 3239m depth on the Davidson seamount, both off the coast of California (Table 1; Figure 1). Regardless of *Osedax* species, *Arcobacteraceae* were the dominant microbial group associated with the worm trunks at early time points (< 24 months, n = 16), constituting 59% of the average recovered Campylobacterales ribotypes, a significantly higher representation than at later time points (18-28%; ANOVA p < 0.02; Figure 5). *Osedax* collected at early time points had relatively few aggregations of bacteria along the trunk that were visible by microscopy, while at later time points bacteria appeared to cover much more of the epithelial surface (Figures 3–4, compared to Figure S4). A specific *Arcobacteraceae* ribotype occurred at the Davidson site, and was related to those associated with other eukaryote hosts (Figure 2). *Sulfurospirillum* appeared to peak in abundance during the mid-stages of whalefall degradation (~50-60 months, n = 9), representing 41% of the recovered Campylobacterales ribotypes, compared to 12-16% at early and late stages (ANOVA p < 0.03; Figure 5). While at later time points (>140 months, n = 12) the dominant genus transitioned significantly to *Sulfurimonas*, averaging 71% of Campylobacterales ribotypes compared to early-mid time periods (25-40%; ANOVA p <0.01; Figure 5). All observed *Sulfurospirillum* and *Sulfurimonas* ribotypes were shared among *Osedax* species, regardless of Davidson or Monterey Canyon sites, suggesting that neither host species nor seafloor location, even at vastly different depths, plays a major role in assembling the *Osedax* Campylobacterales community (only 20-30% of the variation was influenced by either factor, according to one-way ANOSIM; Figure S5).

**Figure 5:**
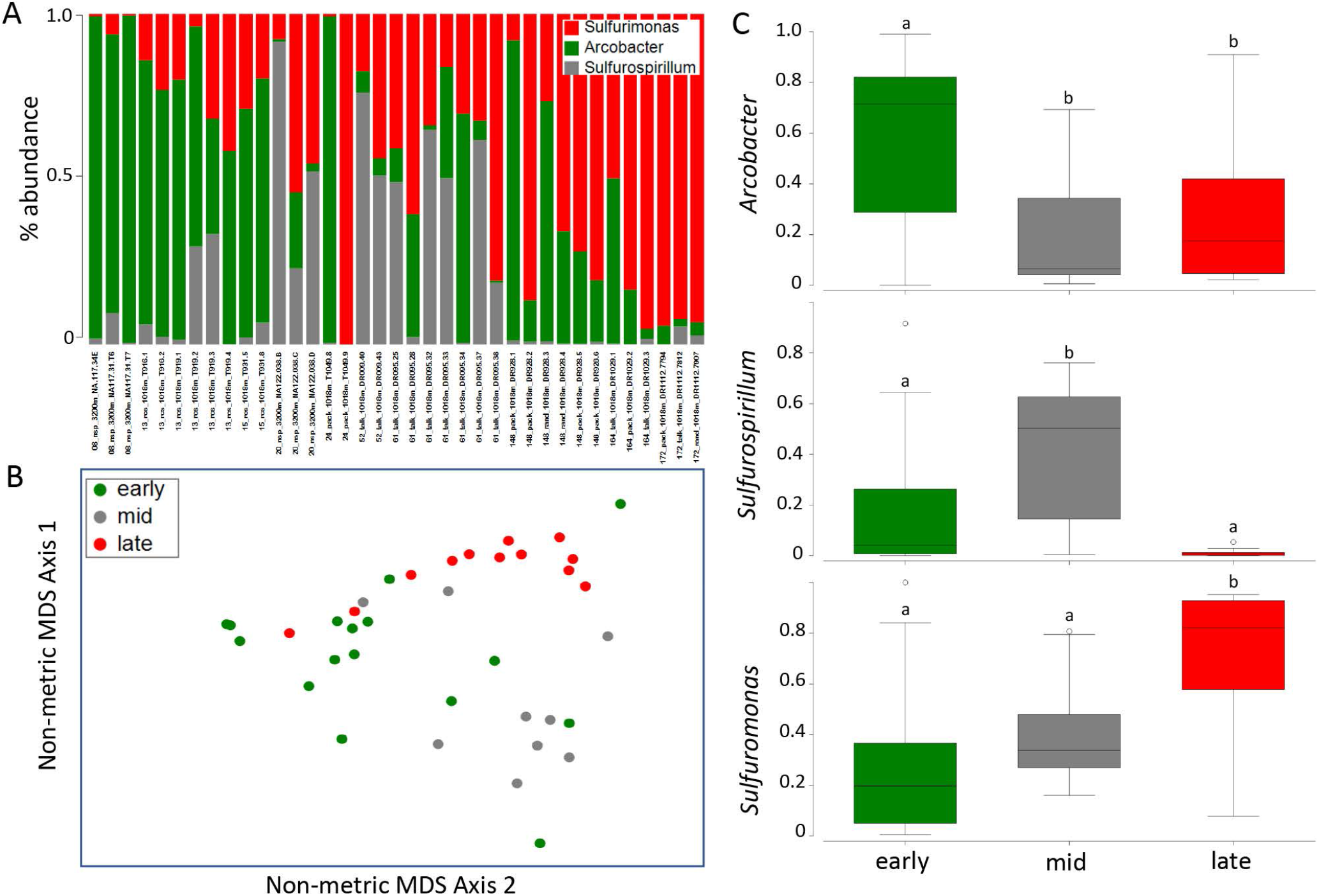
Dominant Campylobacterales 16S rRNA gene sequences recovered from barcoding of 6 *Osedax* species at 2 whalefalls off of northern California, USA. (A) Relative abundance of the genera *Arcobacter, Sufurospirillum* and *Sulfurimonas*, based on 16S rRNA gene amplicon barcoding. (B) Non-metric multidimensional scaling (NMDS) ordination of Campylobacterales communities associated with *Osedax* (square root transformation; Bray-Curtis similarity). Each point represents all Campylobacterales 16S rRNA gene sequences recovered from a single specimen. Ordination comparing 3 different time points from the 1018-m whalefall; early (13-24 months), mid (52-61 months), and late (148-172 months). ANOSIM p < 0.01 for all 3 comparisons, suggesting a significant difference between the timeframes, but with some overlap (R = 0.35-0.53). (C) Box plots of the three dominant *Osedax*-associated genera, as relative percent abundance, at 2 different whalefalls and 3 different time points; early, mid, and late. (n = 16, 9, 12, resp). Levels of significance based on ANOVA p < 0.05. Data points outside of the 25–75% range are identified by open symbols.

To better understand the physiological potential, and therefore ecological influences, of the *Osedax* epibionts, we performed metagenomic sequencing of a single specimen of *Osedax frankpressi* collected from a 3^rd^ whalefall at 2891 m depth in Monterey Canyon. We identified four near complete genomes of the dominant *Osedax* epibionts (completeness scores of 93-100%; 0.6-6.8% contamination; Table 2), with nearly identical 16S rRNA gene sequences to those recovered via barcoding (99.6-100%; Figure 5). These genomes belonged to *Arcobacter* (sensu lato, closely related to *Arcobacter nitrofigilis*, the type species of the genus; 50), *Sulfurospirillum* and *Sulfurimonas*, the 3 dominant Campylobacterales epibionts, the latter represented by 2 distinct genomes identifiable by a large difference in sequencing coverage depth (16 vs 268X; Table 2). The ‘high-coverage’ *Sulfurimonas* (268X), referred to further in the following metagenomic sections unless otherwise noted, was in far greater abundance than even the well-known primary Oceanospirillales endosymbiont (at 30X coverage). A single Kordiimonadales (Alphaproteobacteria) genome was also recovered from the external surface of *Osedax*, however, our microscopy analysis did not indicate integration into the *Osedax* epithelium, so we do not focus on it further (genome available at # PRJNA813420).

| Taxon | coverage | # of<br>contigs | genome<br>size | N50 | rrna<br>copy # | BUSCO<br>Completeness | CheckM |  |
| --- | --- | --- | --- | --- | --- | --- | --- | --- |
|  |  |  |  |  |  |  | Completeness | Contamination |
| *Arcobacter* epibiont | 34x | 1 | 2901687 | 2901687 | 6 | 93.1 | 93.5 | 6.79 |
| *Sulfurospirillum* epibiont | 13x | 33 | 2726073 | 225838 | 2 | 92.7 | 95.1 | 4.27 |
| *Sulfurimonas* epibiont HC | 268x | 1 | 2827517 | 2827517 | 12 | 98.6 | 100 | 0.61 |
| *Sulfurimonas* epibiont LC | 16x | 1 | 2607188 | 2607188 | 12 | 95.1 | 98.7 | 1.02 |

An analysis of genomic motifs involved in metabolism revealed differences in the capabilities among the *Osedax* epibionts. In general, there appeared to be a shift in metabolic strategy from heterotrophy to autotrophy along the temporal successional gradient, from *Arcobacter* (s.l.) and *Sulfurospirillum* at early-mid time points to *Sulfurimonas* at later time points (Figure 6). All of the epibionts can metabolize hydrogen using a shared suite of hydrogenase enzymes (Groups 1, 2a and 4), although gene copy numbers vary. The epibionts differ, however, in their capacity to metabolize oxygen, carbon, nitrogen, and sulfur. For oxygen metabolism, *Arcobacter* and *Sulfurospirillum* possess *cyoE* and the cytochrome oxidase genes *coxA/B* - involved in processing heme and electron transport during aerobic respiration –which are absent in *Sulfurimonas* (Figure 6). All *Osedax*-associated epibionts can reduce nitrate (via *napA/B*), however, only the *Sulfurimonas* epibionts possessed the *nirS* gene for nitrite reduction. In addition, the *Arcobacter* and *Sulfurimonas* possess genes involved in the reduction of nitric/nitrous oxide that are absent in the *Sulfurospirillum*. By contrast, the *Sulfurimonas*, which dominates the trunk surface at later stages of host decomposition, contained genes involved in carbon fixation through the reverse TCA cycle (*aclA* and *aclB*; Figure 6) that are not found in the other epibionts. While all 3 genera can utilize sulfur compounds, the high-coverage *Sulfurimonas* epibiont has additional genes involved in sulfur metabolism, including two copies each of the sulfide:quinone oxidoreductase (*sqr*) and the *soxZ* gene, in addition to the full thiosulfate oxidation (*sox*) pathway. The *sox* pathway was notably absent from the low-coverage *Sulfurimonas* strain (Figure 6).

**Figure 6:**
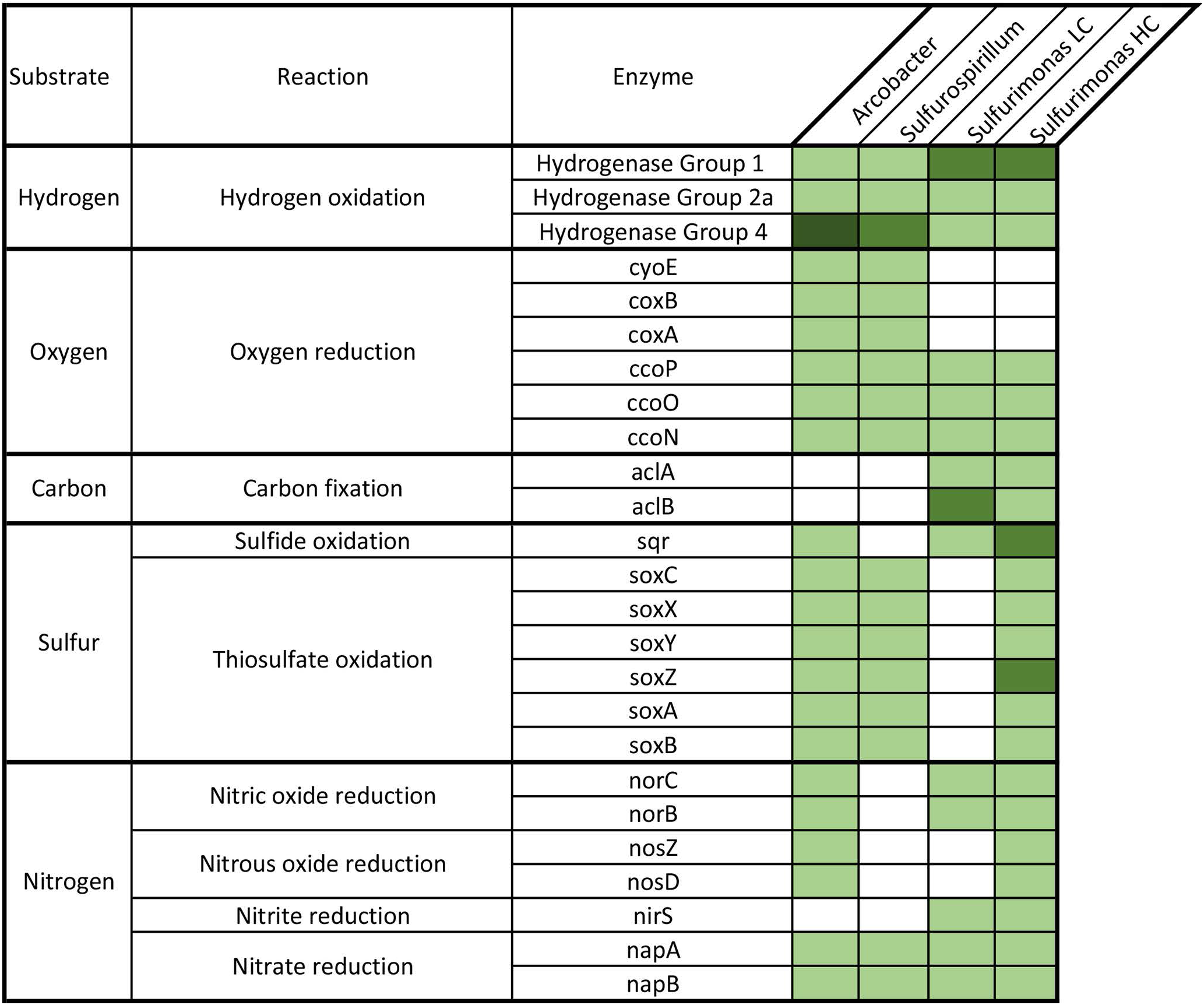
Comparison of genes involved in hydrogen, oxygen, carbon, sulfur, and nitrogen (H,O,C,S,N) metabolism that are present in the Campylobacterales epibionts associated with *Osedax*, using a Hidden Markov Motif (HMM) gene identification analysis through LithoGenie. The number of matches to the metabolic HMMs were quantified for each sample and visualized as a heatmap, including the copy number of each gene. Compared to one another, the epibionts showed a shift in their degrees of investment to carbon and sulfur metabolism later in the degradation process, when *Sulfurimonus* dominates, reflected by their total number of genes involved in each biochemical process.

The average amino acid identity (AAI) of the *Osedax* epibionts differed considerably from close relatives (*N* = 50; 55-78%), suggesting significant divergence time between the epibionts and free-living relatives with available genomes (Table S1). Despite having similar genome sizes to free-living deep-sea lineages, all 3 *Osedax*-associated Campylobacterales have decreased coding densities, and significantly more transposable elements which represented ~2-6% of their genomes, compared to < 1 % for all but a few free-living relatives (Tables 3, S1). A pan-transposase analysis revealed that the epibionts shared numerous insertion sequence families among them; however, they did not share any with the primary symbiont (Table S2). Functional genes carried on transposons included those that encode for a Type I restriction modification system (*hsdM* superfamily), a leukotoxin export ATP-binding protein *ltxB*, and a toxin of the *relE/parE* family, which were shared by all *Osedax* Campylobacterales. An additional membrane fusion protein (MFP) of a Type 1 secretion system (T1SS) was also identified on a transposon in the high-coverage *Sulfurimonas* epibiont (Table S3). Lastly, all *Osedax* Campylobacterales shared a Mu-like bacteriophage, which was not present in free-living relatives, or in the primary Oceanospirillales endosymbiont. The phage appeared intact in the high-coverage *Sulfurimonas*, and is ~19 kb in size, with 15 open reading frames (ORFs), 13 of which code for proteins (7 are known viral proteins, 6 are hypothetical; Table S4), and two insertion sequences (*attL* and *attR*).

| Bacterial ID | Genome size (bp) | # secreted proteins | % of SP with ELP | # insertion sequences | % IS (by length) |
| --- | --- | --- | --- | --- | --- |
| ***Arcobacter* epibiont** | **2901687** | **349** | **59.3** | **227** | **4.5** |
| *Arcobacter* anaerophilus (CP041070_0) | 3016922 | 159 | 13.2 | 10 | 0.6 |
| *Arcobacter* aquamarinus (CP042812_0) | 2829476 | 175 | 8.0 | 8 | 0.5 |
| *Arcobacter* butzleri (CP000361_0) | 2341251 | 146 | 2.7 | 3 | 0.4 |
| *Pseudoarcobacter* acticola (CP042652_0) | 3019071 | 256 | 36.7 | 105 | 4.2 |
| GCA_000585115.1 | 2287768 | 185 | 6.5 | 7 | 0.8 |
| GCA_000585155.1 | 2496885 | 178 | 12.9 | 27 | 1.3 |
| ***Sulfurospirillum* epibiont** | **2726105** | **313** | **41.5** | **205** | **6.5** |
| *Sulfurospirillum* deleyianum (GCA_000024885) | 2306351 | 97 | 32.0 | 35 | 1.7 |
| *Sulfurospirillum* multivorans (GCA_000568815) | 3175729 | 204 | 48.0 | 73 | 2.4 |
| GCA_000265295.1 | 2510109 | 90 | 23.3 | 13 | 0.9 |
| GCA_002205395.1 | 2876607 | 176 | 41.5 | 8 | 0.4 |
| GCA_002309535.1 | 2814086 | 148 | 35.1 | 20 | 0.7 |
| GCA_008083195.1 | 3181530 | 202 | 47.5 | 67 | 2.3 |
| ***Sulfurimonas* epibiont (HC)** | **2827517** | **255** | **46.3** | **68** | **2.5** |
| ***Sulfurimonas* epibiont (LC)** | **2607188** | **196** | **30.6** | **41** | **2.2** |
| *Sulfurimonas* autotrophica (CP002205) | 2153198 | 112 | 20.5 | 11 | 1.1 |
| *Sulfurimonas* denitrificans (CP000153) | 2201561 | 112 | 13.4 | 8 | 0.9 |
| *Sulfurimonas* sediminis (CP041235_0) | 2320257 | 154 | 39 | 76 | 2.9 |
| GCA_000242915.2 | 2952682 | 321 | 62 | 24 | 0.7 |
| GCA_000445475.1 | 2302023 | 106 | 17 | 9 | 0.6 |
| GCA_009192995.1 | 2093483 | 106 | 14.2 | 6 | 0.8 |

The *Osedax*-associated *Arcobacter* and *Sulfurimonas* also possessed additional genes that encode proteins involved in attachment and secretion system machinery (Table S3). For example, the *Arcobacter* contained 5 copies of the Type 5a secretion system (T5aSS), compared to 0-2 copies in close relatives. The *Sulfurospirillum* epibiont had a tight adherence (TAD) pilus, distinguishing it from close relatives. Similarly, the high-coverage *Sulfurimonas* epibiont had a Type 5a secretion system absent in close relatives, and a complete 14-gene Type 6 secretion system (T6SS) containing 15 copies of the *tssD* gene and 21 copies of the *tssI* gene — The *tssD* and *tssI* genes encode 2 of the 12 core T6SS subunits; the stacked hexameric rings (i.e., Hcp tube) that extend outward from the bacterial cell membrane and the distal cell-puncturing device (a trimer of VgrG), respectively.

Finally, the *Osedax* Campylobacterales genomes contained hundreds of genes encoding predicted secreted proteins, and within these, significantly more (31-59%) had eukaryotic-like protein domains (ELPs) compared to free-living relatives (2-30X; Tables 3, S1). These ELPs, which can be mobilized by the secretion systems described above, comprised 54 families, based on Pfam identification, plus 7 others of unknown function (Table S5). Several of the ELPs were shared among the 3 main *Osedax* Campylobacterales, including: ATP:guanido phosphotransferase (N-terminal domain), an integrase core domain, and a Helix-turn-helix (HTH)-like domain. The early colonizers *Arcobacter* and *Sulfurospirillum* also shared a homeobox-like domain, coding proteins in a large family of transcription factors that contain a highly conserved DNA-binding domain, and a second integrase core domain. The high- and low-coverage *Sulfurimonas* genomes, not unexpectedly based on their shared evolutionary history, shared many of their ELPs (~50%; Table S5).

## Discussion

Over the course of 14 years, we identified pervasive Campylobacterales epibionts associated with the external surface of 7 *Osedax* species from deep-sea whalefalls off of northern California (1018-3239m depth). The persistence of this specific bacterial order, which had been observed with *Osedax* previously (2, 10, 11), supports the assertion by Verna et al. (2010) that this relationship is more than transitory. Metagenome analysis suggests a long-term association between the Campylobacterales and their *Osedax* hosts based on an abundance of genes encoding secretions systems that are absent in free-living relatives, perhaps to ensure attachment to the host, and an enrichment in secreted proteins with eukaryotic-like protein (ELP) domains. ELPs, which can be rare for some groups (ex. *Arcobacteraceae*), are considered a bacterial strategy for modulating eukaryotic processes and, in a few symbiotic systems, have been implicated in extracellular secretion, cell binding, colonization, and protein-protein interactions (60–62). Some of these domains may even encode signal peptides that interact with secretion systems (63), which were also observed in the *Osedax*-associated Campylobacterales epibionts. ELPs can either be acquired by horizontal gene transfer from a eukaryote, followed by divergent evolution, or through convergent evolution with a non-homologous protein, both of which would take time to evolve. Additionally, they also shared a Mu-like bacteriophage carrying numerous unknown genes. The pronounced abundance of mobile elements in the *Osedax*-associated Campylobacterales, including toxin genes shared between them, suggests a dynamic transfer of genetic material between the microbes, either via cell-to-cell contact or phage transfer.

The relationship between *Osedax* and the Campylobacterales is not fixed, as the trunk epidermis is repeatedly exposed and re-colonized throughout the course of whalefall degradation. Temporal succession has also been observed for the *Osedax* host species (4) and their primary symbionts (7–8), as well as free-living microbial communities in the surrounding sediments (9) as the whale carcass degrades, suggesting a direct influence of the local environment on associated microbial populations (7). On the trunk epidermis of *Osedax, Arcobacteraceae* was the dominant founding group, despite differences in water depth, seafloor location, and host species. Similar species-level ribotypes were also recovered from *O. mucofloris* collected from a Minke whalefall off the coast of Sweden, at 36 months post implantation (11, 66), and from *O. roseus* collected at 3 months from the 1018 m depth in Monterey Canyon sampled in this study (10). The *Arcobacteraceae* is a known early colonizer in sulfur-rich habitats (67–68). They have also been identified as the pioneer producer of floc during in situ and shipboard experiments with bacterial biomass collected from hydrothermal vents at 9°N East Pacific Rise (69). Additionally, a recent study demonstrated that the microbial community composition of *Arcobacter, Sulfurimonas*, and *Sulfurovum* in hydrothermal vent fluid incubations was highly dependent on oxygen levels (70), a parameter that also varies dramatically at whalefalls (71). A general predisposition for oxic environments by the *Osedax*-associated *Arcobacter* is indicated by the possession of the heme O synthase gene *cyoE* and the cytochrome oxidase genes *coxA/B*, involved in electron transport during aerobic respiration. The *Osedax*-associated *Arcobacter* (s.l). is a heterotroph, likely dependent on organic substrates available on the host surface. Similar to the *Arcobacter* recovered from *Lenisia*, a Breviatea protist, the *Osedax*-associated *Arcobacter* possesses numerous hydrogenase genes and those involved in a cellular response to nitric/nitrous oxide, which underpin the mutual benefits in the *Lenisia*-*Arcobacter* symbiosis through the transfer of hydrogen (52). The relevance of this capability as a symbiont specificity determinant, as observed in other symbioses (72), remains unconfirmed for both the *Osedax*-associated *Arcobacter* and *Sulfurimonas*.

*Sulfurospirillum* was associated with *Osedax* trunk surfaces throughout the duration of this study, but was most prominent during the intermediate time frames from ~50-60 months, suggesting some adaptability during the transitional stages of organic carbon breakdown. This genus is globally found in deep-sea habitats rich in sulfur compounds (73) and the *Osedax-* associated phylotype was similar in 16S rRNA gene to those found in whalefall environments (2, 19). The only other host-associated *Sulfurospirillum* described thus far is a heterotrophic, hydrogen-utilizing epibiont of the vent worm *Alvinella pompejana* (77). Unlike the *A. pompejana* epibiont, the *Osedax*-associated *Sulfurospirillum* is missing the *phsA* and *sqr* genes, so must rely on exogenous thiosulfate (77). This may explain why it never dominated the *Osedax* trunk community by itself, but rather co-occurred with either *Arcobacter* or *Sulfurimonas*, both of which can oxidize sulfide to thiosulfate. Additionally, the genome of the *Osedax*-associated *Sulfurospirillum*, unlike close relatives, possesses a gene cluster encoding a tight adherence (TAD) pilus, the adhesive structure often used for colonization of surfaces, including eukaryote hosts (78–79).

The chemoautotrophic *Sulfurimonas* dominated the epibiont community associated with *Osedax* at later stages of whale decomposition (> 140 months). Via metagenomic analysis, two *Sulfurimonas* genomes were recovered, one of which was far more abundant than even the primary intracellular symbiont, based on a ten-fold higher genome coverage depth. The hydrogen and sulfur-utilizing capabilities of the autotrophic *Osedax*-associated high coverage *Sulfurimonas* appears to be similar to two other Campylobacteria, *S. paralvinellae* and *S. hydrogeniphila*, isolated from a deep-sea tubeworm ‘nest’ and hydrothermal vent chimney, respectively (55, 81). A community shift from *Arcobacter* and *Sulfurispirillum* (both generally organotrophs) to autotrophic *Sulfurimonas* species is likely influenced by changes in the chemical environment of decomposing organic matter. Kalenitchenko et al. (2016) noted a temporal transition from chemoorganotrophic metabolism to chemoautotrophic reliance in reduced deep-sea wood mesocosms (68). A similar shift from *Arcobacter* to *Sulfurimonas* at hydrothermal ‘snowblower floc’ eruptions has been attributed to elevated hydrogen sulfide levels, and the subsequent utility of both the *sox* and *sqr* systems by *Sulfurimonas* (82). Indeed, the high-coverage *Osedax*-associated *Sulfurimonas* genome possessed type II and type IV sulfide:quinone oxidoreductase genes, which encode a key enzyme involved in sulfide homeostasis (oxidation and assimilation) and detoxification in bacteria (81), which may help protect its host from harmful by-products during late stages of whale decomposition. Dominance of the high coverage *Sulfurimonas* at latter stages of whale carcass decomposition is likely due to the numerous reduced sulfur compounds the bacteria can use as an energy source (76), and the deployment of the epibionts arsenal of secretion systems, that are often used to form biofilms and to gain a competitive advantage over neighboring bacteria, as observed in both the *Euprymna* squid and legume symbioses (83–84).

With the exception of obvious nutritional episymbioses, such as ciliates, nematodes and yeti crabs (85–89), the role of attached external bacteria in supporting host health has received relatively little attention. With regard to *Osedax*, Borchert et al. (2021) proposed that the host could benefit nutritionally from enhanced dissolution of inorganic bone components by proton release and subsequent acidification by the sulfur-oxidizing epibionts (80). This appears unlikely based on the distinct lack of epibiotic bacteria on the root surfaces, the only tissue in contact with the bone. Our analysis suggests that detoxification of sulfide by the bacteria may be a possible benefit to the *Osedax* worm host. Hydrogen sulfide is likely to emanate from the whale carcass, especially at later stages of decomposition, and thus sulfide-oxidizing bacteria positioned near the tissue-bone interface could convert this sulfide to less toxic products. It is generally accepted that bacteria associated with epithelial surfaces produce metabolic by-products that can be absorbed across the epithelial barrier, thereby influencing the host (90), however whether the Campylobacterales bacteria of *Osedax* are commensals or beneficial remains undetermined.

## Conclusion

Ecological factors shaping the epibiont communities of marine organisms remain poorly understood. The recurrence of three Campylobacterales associated with diverse *Osedax* species collected from multiple deep-sea locations suggests they are specific epibionts that share a long-evolutionary history with their host. All three epibionts have an affinity for organic-rich and sulfide-rich habitats, however a successional shift in their composition reveals that they are a dynamic community that changes over time. Factors shaping the epibiome may include the metabolic capabilities of the bacteria themselves, host-controlled changes to the epidermis, and the chemically-diverse abiotic conditions (ex. sulfides, oxygen and nutrients) that change as the whale carcass habitat degrades over time. Our metagenomic analysis revealed the *Osedax*-associated Campylobacterales to possess genes that allow them to both fuse with the host epithelium, and subsequently take advantage of the metabolic opportunities in their changing environment while attached to a host. The presence of extensive secretion systems may also influence their composition, by moderating interactions with *Osedax* and/or competing microbes. Our results provide evidence of a persistent yet dynamic relationship between *Osedax* and specific Campylobacterales epibionts that possess unique genomic features. However, the role of the biofilm on the physiology of *Osedax* remains unknown.

## Methods

### Specimen Collection

*Osedax* specimens were collected from a whalefall at 1018m depth in the Monterey Canyon off the coast of California (from 2005-2019), using the remotely operated vehicles (ROVs) Tiburon or Doc Ricketts (on the R/V Western Flyer), and from a whalefall at 3239m depth on the Davidson seamount (from 2019-2020), using the ROV Hercules (on the R/V Nautilus; Table 1). The whalefall at 1018m in the Monterey Canyon (36.772°N/122.083°W) was implanted by the Monterey Bay Aquarium Research Institute in October 2004 (ref. 4). The whalefall on the Davidson seamount (35.582°N/122.629°W) was discovered serendipitously in October 2019. Several additional specimens of *O. frankpressi*, used for metagenomic analysis and microscopy, were collected from a natural whalefall in Monterey Canyon at 2891 m (36.613°N/ 122.434°W). At the 1018-m site, *Osedax* worms were collected between 8-172 months after the carcass was first deposited on the seafloor, or discovered in an early stage of decomposition (Table 1). Whalefall stages were categorized as being in early, mid, or late stages by the progression of bone degradation; ‘early’ designated as having significant whale tissue and bone biomass present; ‘mid’ designated as having little whale tissue present, and ‘late’ stages designated by extreme reduction in bone biomass (Figure 1). Pursuant to the Marine Mammal Protection Act (50 CFR 216.22 and 216.37), authorization was received for the Monterey Bay Aquarium Research Institute and the Monterey Bay National Marine Sanctuary (MBNMS-2020-006) to collect whalefall specimens for scientific purposes during exploratory dives via remotely-operated vehicles in the Monterey Bay National Marine Sanctuary. Additionally, a general CDFW collecting permit SC-10578 (S. Goffredi) was acquired for collection of *Osedax* specifically. All *Osedax* species used in this study have been previously described, with the exception of one undescribed species from the Davidson Seamount from dive H1796 (Table 1).

### Microscopy

Specimens for fluorescence in situ hybridization (FISH) microscopy were initially preserved in 4% sucrose-buffered paraformaldehyde (PFA) and kept at 4°C. These PFA-preserved specimens were rinsed with 2× PBS, transferred to 70% ethanol, and stored at −20°C. Tissues were dissected and embedded in Steedman’s wax (1 part cetyl alcohol: 9 parts polyethylene glycol (400) distearate, mixed at 60°C). An ethanol: wax gradient of 3:1, 2:1 and 1:1, and eventually 100% wax, was used to embed the samples (1 h each treatment). Embedded samples were sectioned at 2-5 μm thickness using a Leica RM2125 microtome and placed on Superfrost Plus slides. Sections were dewaxed in 100% ethanol rinses. The hybridization buffer included 35% formamide, fluorescent probes were at a final concentration of 5 μg/ml, and the wash solution contained 450 mM NaCl (10). Initially, we used the epsilonproteobacteria-specific probe EPS549; ref. 23) labelled with FITC or Cy3. A universal bacterial probe (Eub338-I; ref. 24), labelled with Cy3, Cy5, or Alexa488, was also used. Finally, two group specific probes were designed (using Geneious Prime 2021; www.geneious.com) for both the *Arcobacter* (Epsi41; 5’-TTAACCCGCCTACATGCTCT-3’) and *Sulfurimonas* (Epsi126; 5’-AATTCCATCTCCC-CCTCCCA-3’), both labelled with Cy3. Probes were hybridized at 46°C for 4-8 h, followed by a 15 min wash at 48°C. Sections were counterstained with 4’6’-diamidino-2-phenylindole (DAPI, 5 mg/ml) for 1 min, rinsed and mounted in Citifluor and examined by epifluorescence microscopy using a Nikon E80i epifluorescence microscope with a Nikon DS-Qi1Mc high-sensitivity monochrome digital camera.

For examination by transmission electron microscopy, samples (approximately 1 mm^3^) were fixed in 3% glutaraldehyde buffered with 0.1 M phosphate and 0.3 M sucrose (pH 7.8). Following a wash in 0.1 M sodium cacodylate containing 24% sucrose, samples were postfixed with 1% OsO4 in 0.1 M sodium cacodylate for 1 h, stained *en bloc* in 3% uranyl acetate in 0.1 M sodium acetate buffer for 1 h, dehydrated through an ethanol series, then infiltrated and embedded in Spurr’s resin (Ted Pella, Redding, CA, USA). Thin (70 nm) sections were stained with methylene blue and lead citrate, respectively, then examined and photographed using a Zeiss Labrolux 12 light microscope and Zeiss EM109 TEM.

### Molecular Analysis

Specimens for molecular analysis were either frozen at −80°C or preserved immediately upon collection in ~90% ethanol. Total genomic DNA was extracted using the Qiagen DNeasy kit (Qiagen, Valencia, CA, USA) according to the manufacturer’s instructions. For some specimens, DNA was extracted from *Osedax* trunk tissues only, while for others, as in small species like *O. talkovici*, whole specimens were used. Select mucous tubes were also extracted, separate from animal tissue.

*Osedax* identity was confirmed by sequencing the mitochondrial cytochrome c oxidase subunit I gene (COI). This gene was amplified using the previously published primers LCO1490/ HCO2198, according to ref 25. Amplification products were sequenced directly using Sanger sequencing, via Laragen Inc. (Culver City, CA, USA), and compared to published sequences in GenBank and in ref. 6.

To characterize the *Osedax*-associated bacterial diversity, we performed 16S rRNA gene amplicon sequencing. The V4-V5 region of the 16S rRNA gene was amplified using bacterial primers with Illumina (San Diego, CA, USA) adapters on the 5’ ends of 515F/806R (ref. 26), with Q5 Hot Start High-Fidelity 2x Master Mix (New England Biolabs, Ipswich, MA, USA) and annealing conditions of 54°C for 25 cycles. Each product (2.5 μl) was barcoded with Illumina NexteraXT index 2 Primers that include unique 8-bp barcodes (64°C annealing temperature and 11 cycles). Secondary amplification products were purified via vacuum manifold (Millipore-Sigma MultiScreen plates (St. Louis, MO, USA) and quantified using QuantIT PicoGreen dsDNA (Thermo-Fisher Scientific; Waltham, MA, USA) on a BioRad CFX96 Touch Real-Time PCR Detection System. Barcoded samples were combined in equimolar amounts (~ 100 ng) into a single tube and purified with the Promega Wizard SV Gel and PCR Clean-Up kit (#A9281) before submission to Laragen (Culver City, CA, USA) for 2 x 300bp paired end analysis on the Illumina MiSeq platform with PhiX addition of 15-20%. MiSeq 16S rRNA gene sequence data was processed in Quantitative Insights Into Microbial Ecology (v1.8.0; ref. 27), using the default parameters. Sequences were clustered into *de novo* operational taxonomic units (OTUs) with 99% similarity using UCLUST open reference clustering protocol, and then, the most abundant sequence was chosen as representative for each *de novo* OTU. Taxonomic identification for each representative sequence was assigned using the Silva-138 database (28), clustered at 99% similarity. A threshold filter was used to remove any OTU that occurred below 0.01% in the combined samples dataset. Analyses are based on Bray-Curtis distances of fourth-root transformed data. Quantification and statistical analyses are described in the Results sections and figure legends.

### Microbial genomes: DNA extraction, sequencing, and bioinformatic analysis

High molecular weight genomic DNA was extracted from an entire *Osedax frankpressi* adult female following the Bionano genomics IrysPrep agar-based animal tissue protocol (Catalogue # 80002). Sequencing of the gDNA was performed at UC Berkeley with a PacBio Sequel II machine to generate long reads of high contiguity, and on an Illumina HiSeq6000 for short reads of high accuracy and depth of coverage. The reads were profiled taxonomically using Kraken (29), to filter out eukaryotic reads, then all prokaryotic reads were co-assembled using MetaFlye (30) with automatic genome size selection followed by 10 polishing iterations. The assembly graphs were manually inspected using Bandage (31) and were binned using MaxBin2 (32) with minimun contig length of 1000 base-pairs maximum iteration of 50, and a probability threshold of 0.9. Each bacterial genome was then polished with both the short and long reads using NextPolish (33) following the recommended configuration. To resolve contamination issues due to heterogeneity of *Sulfurospirillum* strains within our sample, we assembled using the short Illumina reads using SPAdes v.3.15.4 (ref. 34) and used BlobTools v.1.1.1 (ref. 35) to select for the most abundant *Sulfurospirillum* strain. *Sulfurospirillum* contigs based on Illumina data were then mapped and polished to long-reads using NextPolish (33), yielding a genome of high contiguity and completeness with low heterogeneity. Assembly metrics were generated using MetaQuast (36), genome completeness and contamination were checked using BUSCO (37) and CheckM (38) and their taxonomic IDs were identified using GTDB-Tk via wgANI (39) and taxonomic placement of the genomes alongside the thousands of references in the GTDB database.

The genomes were annotated using Prokka (40) using --kingdom Bacteria --gcode 11 -- compliant, and amino acid translations of the annotations were used in OrthoVenn2 (41) for gene enrichment analysis following default parameters. Hidden Markov Motifs (HMMs) involved in metabolism were identified using Lithogenie through the MagicLamp tool (42) utilizing the curated enzymatic motifs from K. Anantharaman (https://github.com/kanantharaman/metabolichmms). Insertion sequences were detected using ISSAGA (43) and IslandViewer4 (44), for all predicted transposable elements and the associated mobilized functional genes. Secreted proteins with eukaryotic like domains were identified using EffectiveELD through EffectiveDB, on default settings (45). Secretion system proteins were annotated using TXSScan (46) through MacSyFinder using curated motifs to check for the presence of genes involved in protein secretion machinery, irrespective of order, to allow the detection of horizontally acquired genes and enriched copies of certain parts of the secretion machinery. Bacteriophages were investigated using Phaster (47) using the genome assembly of each epibiont as fasta input. One sample T tests to report on the significant differences in genomic features between the epibionts and free-living relatives were conducted using ggpubr v.0.1 (ref. 48). The absence of carbon, sulfur and nitrogen metabolism genes in *Sulfurospirillum-related* reads was confirmed by screening the unbinned reads.

## Acknowledgements

The research expeditions were made possible via support from the Monterey Bay Aquarium Research Institute and the Monterey Bay National Marine Sanctuary. We thank the captain and crew of the R/V *Western Flyer*, the pilots of the ROVs Tiburon and Doc Ricketts, the captain and crew of the E/V *Nautilus* and pilots of the ROV Hercules, R.C. Vrijenhoek for expedition leadership, S Johnson for support at sea and in the laboratory, S. Connon for assistance with microbial community analysis. In addition, undergraduates T. Talbert and R. Marin assisted with microscopy analysis, sponsored by the Occidental College Undergraduate Research Center and Ron and Susan Hahn. Support for S.K.G. was provided, in part, by a National Science Foundation grant (IOS-0923775), by NERC IRF (NE/M018016/1) to L.M.H, Wellcome Trust Seed Award (213981/Z/18/Z) to J.M.M.D.

## Data Availability

The raw Illumina 16S rRNA gene barcode sequences and metadata collected in this study are available from the NCBI Small Read Archive (project number PRJNA813533). The 4 *Osedax-* specific Campylobacterales genome assemblies, as well as an Alphaproteobacteria, were deposited to Genbank under project number PRJNA813420.

